# The outer mitochondrial membrane protein TMEM11 is a novel negative regulator of BNIP3/BNIP3L-dependent receptor-mediated mitophagy

**DOI:** 10.1101/2022.03.29.486240

**Authors:** Mehmet Oguz Gok, Jonathan R. Friedman

## Abstract

Mitochondria play critical roles in cellular metabolism and to maintain their integrity, they are regulated by several quality control pathways, including mitophagy. During BNIP3/BNIP3L-dependent receptor-mediated mitophagy, mitochondria are selectively degraded by the direct recruitment of the autophagosome biogenesis protein LC3. BNIP3 and/or BNIP3L are upregulated situationally, for example during hypoxia and developmentally during erythrocyte maturation. However, it is not well understood how they are regulated at steady-state. Here, we find that the poorly characterized mitochondrial cristae morphology regulator TMEM11 unexpectedly localizes to the outer membrane where it forms a complex with BNIP3 and BNIP3L. Loss of TMEM11 causes mitochondrial morphology defects in a BNIP3/BNIP3L-dependent manner and, further, we find that mitophagy is hyper-active in the absence of TMEM11 during both normoxia and hypoxia. Our results reveal a non-canonical role for TMEM11 as a negative regulator of BNIP3/BNIP3L-mediated mitophagy and suggest that the TMEM11/BNIP3/BNIP3L complex coordinately regulates mitochondrial quality control.

## Introduction

Mitochondria play fundamental roles in many cellular processes, including energy production, and are hubs of cellular metabolism. In order to effectively perform these jobs, mitochondria are organized into an elaborate tubular network that is distributed throughout the cell. Mitochondria are enclosed by two membrane bilayers and the inner mitochondrial membrane (IMM) forms elaborate cristae invaginations that compartmentalize the process of oxidative phosphorylation (Pfanner et al., 2019). Several interrelated processes contribute to the spatial organization of mitochondria, including dynamic movements along the cytoskeleton, division and fusion of the organelle, cristae shaping and organizing proteins inside mitochondria, and quality control mechanisms that ensure functional integrity of the organelle. However, we still lack a complete mechanistic understanding of each of these individual processes and how they are coordinated to contribute to the homeostasis of the mitochondrial network.

Mitochondria respiratory function is maintained by the physical organization of cristae membranes inside the organelle. While several determinants shape cristae, the chief organizer of internal organization is the Mitochondrial Contact Site and Cristae Organizing System (MICOS) complex, which enriches at cristae junctions, sites of cristae invagination (Colina-Tenorio et al., 2020). MICOS is comprised of seven core subunits in human cells and associates in a mega-complex, termed the Mitochondrial Bridging Complex (MIB), with proteins on the outer mitochondrial membrane (OMM), including the beta-barrel assembly SAM complex (Ott et al., 2012). MICOS/MIB also physically interacts with mitochondrial network shaping proteins such as the mitochondrial motility factor Miro (Li et al., 2021; Modi et al., 2019). Thus, MICOS/MIB spans the intermembrane space and is capable of coordinating mitochondrial internal organization with external determinants.

Mitochondrial function depends not only on the internal shape of cristae membranes, but also on processes that maintain the overall performance of the network. Several quality control processes deal with insults such as inappropriate protein targeting or unfolded proteins (Ng et al., 2021). During severe stress that causes the inability of mitochondria to maintain membrane potential, the PINK/PARKIN pathway can recruit the ubiquitin-proteosome system to mediate turnover of the organelle via mitophagy. However, basal mitophagy has been observed *in vivo* independent of the PINK/PARKIN pathway (Lee et al., 2018; McWilliams et al., 2018). Alternative pathways include receptor-mediated mitophagy, whereby outer mitochondrial membrane proteins can selectively recruit the autophagosome biogenesis protein LC3 through cytosolically-exposed LC3-interacting motifs (LIR domains) (Liu et al., 2014). The best characterized of these mitophagy receptors are the BCL2 family members BNIP3 and BNIP3L, which are upregulated to deal with stress insults such as hypoxia and mediate mitophagy during developmental processes (Bellot et al., 2009; Moriyama et al., 2014; Novak et al., 2010; Ordureau et al., 2021; Sandoval et al., 2008; Schweers et al., 2007; Simpson et al., 2021; Zhang et al., 2008). However, these proteins can frequently be detected at lower levels on mitochondria prior to such upregulation (Bellot et al., 2009; Glick et al., 2012; Ordureau et al., 2021), raising the question of how these proteins are regulated at steady-state.

Previously, the IMM protein TMEM11 has been associated in the process of cristae organization, though contributes to mitochondrial morphology through an unknown functional role. Depletion of TMEM11 in human cells and mutations in the *Drosophila* homolog of TMEM11 (PMI) cause severe mitochondrial morphology defects, including mitochondrial enlargement and aberrantly elongated cristae (Macchi et al., 2013; Rival et al., 2011). These mitochondrial morphology defects correspond to whole animal physiological defects, and TMEM11/PMI mutant flies have motor neuron defects and reduced lifespan (Macchi et al., 2013). While high throughput yeast two-hybrid interactome data originally implicated TMEM11 as a BNIP3/BNIP3L interactor (Rual et al., 2005), these data are inconsistent with its previously characterized localization to the IMM (Rival et al., 2011). More recently, proteomic analysis of several MICOS components commonly identified TMEM11 as a MICOS interactor and putative auxiliary subunit (Guarani et al., 2015), which is consistent with the mitochondrial morphology defects that occur in its absence.

Here, we explore the functional role of TMEM11 and its contribution to mitochondrial morphology and function in human cells. We find that while TMEM11 associates with the MICOS complex, it localizes to the OMM where it directly interacts and stably forms a complex with the mitophagy receptors BNIP3 and BNIP3L. Further, we find that that BNIP3 and BNIP3L are primarily responsible for the mitochondrial morphology defects of TMEM11-depleted cells. Finally, we show that loss of TMEM11 sensitizes cells to both basal and hypoxic mitophagy mediated by BNIP3/BNIP3L. Thus, TMEM11 is a novel negative regulator of the mitophagy receptors BNIP3/BNIP3L.

## Results

### TMEM11 is required for the maintenance of normal mitochondrial morphology

To ascertain the functional role of TMEM11, we utilized CRISPR interference (CRISPRi) (Qi et al., 2013) to stably deplete TMEM11 from U2OS cells. Cells constitutively expressing the transcriptional repressor dCas9-KRAB (Le Vasseur et al., 2021) were transduced with an integrating lentiviral plasmid co-expressing TagBFP and either a scrambled control sgRNA or sgRNAs targeting the transcription start site of TMEM11, and TagBFP-positive cells were selected by FACS. Two different stable knockdown lines were generated that exhibited nearly complete depletion of TMEM11 as assayed by Western analysis (Fig. 1A). We then stained cells with the vital dye Mitotracker and examined mitochondrial morphology by fluorescence microscopy (Fig. 1B). Consistent with previous work (Rival et al., 2011), more than half of the cells in each TMEM11-depleted cell line exhibited mitochondria that were enlarged and/or bulbous as compared to the narrow tubular mitochondria observed in control cells (Fig. 1B-1C).

**Figure 1.**
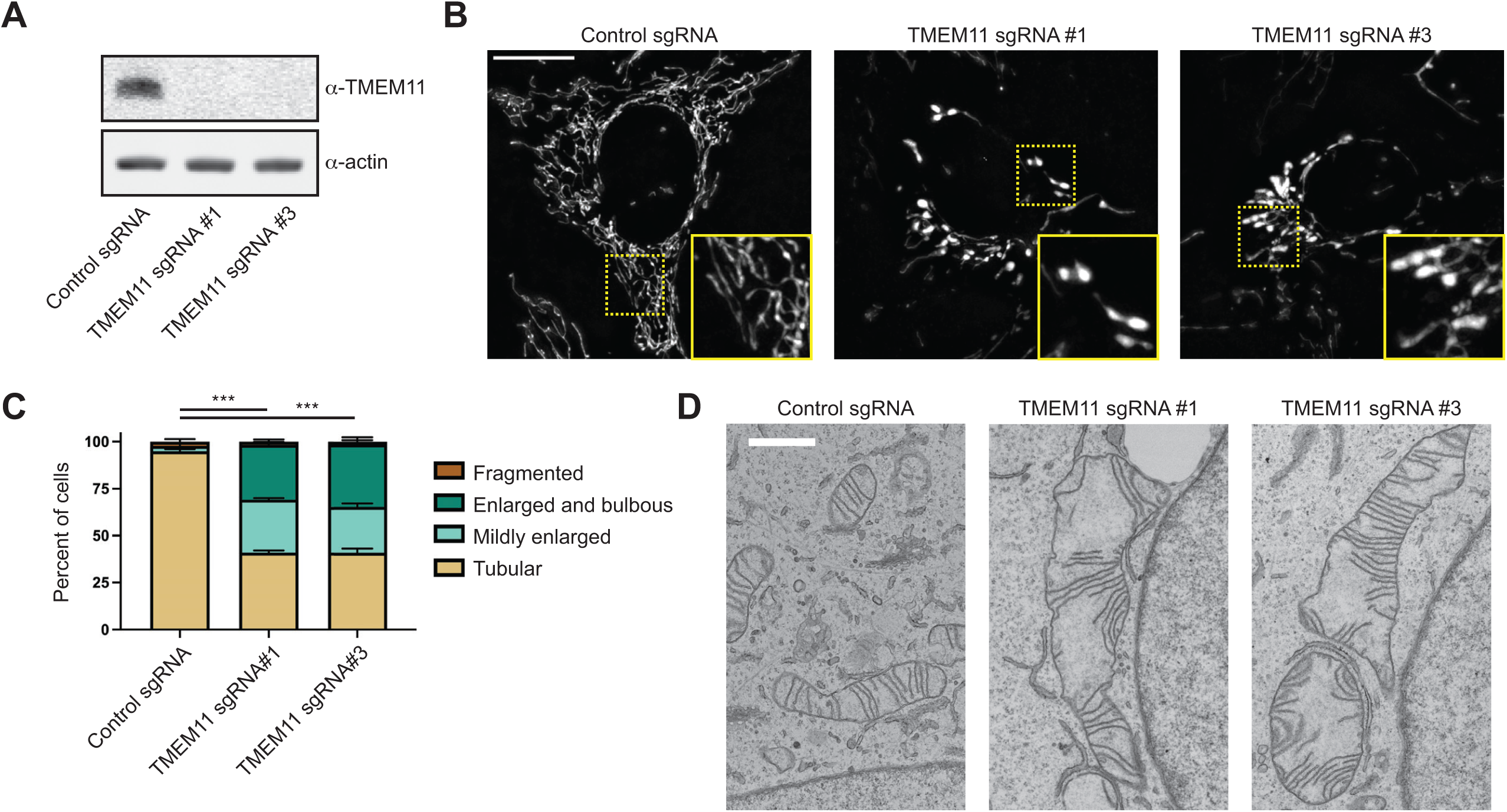
TMEM11 is required for maintenance of normal mitochondrial morphology. **(A)** Western blot analysis of whole cell lysates from U2OS CRISPRi cells expressing scrambled control sgRNA or sgRNAs targeting TMEM11 and probed with the indicated antibodies. **(B)** Deconvolved maximum intensity projections of fluorescence microscopy images are shown of U2OS CRISPRi cells stably expressing the indicated sgRNAs and stained with the mitochondrial dye Mitotracker Deep Red. Insets correspond to dotted boxes. Scale bar = 15 µm. **(C)** A graph of the categorization of mitochondrial morphology from cells as in (B). Data shown represent approximately 100 cells per condition in each of three independent experiments and bars indicate S.E.M. Asterisks (***p<0.001) represent unpaired two-tailed *t* test. **(D)** Representative electron micrographs of mitochondria from CRISPRi cells expressing the indicated sgRNA. Scale bar = 1 µm.

Consistent with its function in human cells, the *Drosophila* TMEM11 homolog PMI is required for the maintenance of normal mitochondrial morphology in flies (Rival et al., 2011). Electron microscopy (EM) analysis of adult brain neuron cell bodies and adult flight muscle from *PMI* mutant flies revealed that mitochondria were enlarged and exhibited elongated and curved cristae membranes compared to those of wild type flies (Macchi et al., 2013). We therefore decided to examine cristae morphology in TMEM11-depleted cells by EM. Strikingly, and consistent with our fluorescence microscopy analysis and the previous EM of *PMI* mutant flies, mitochondria were frequently enlarged in TMEM11-depleted cells (Fig. 1D). Cristae membranes also were curved and/or highly elongated, frequently spanning the width of the enlarged mitochondria (Fig. 1D). These data suggest that TMEM11 contributes to mitochondrial morphology in a conserved manner.

### TMEM11 interacts with MICOS but is not required for the stability or assembly of the complex

Previous proteomic analysis of multiple MICOS subunits identified TMEM11 as a common interacting protein (Guarani et al., 2015). Based on the observed defects of cristae organization in TMEM11-depleted cells (Fig. 1D) and its localization to the IMM (Rival et al., 2011) where it interacts with the MICOS complex, we considered the possibility that TMEM11 associates with the assembled MICOS complex. In mammalian cells, because MICOS subunits assemble into a core complex as well as the MIB complex, its subunits compose complexes that range in size between ∼700 kDa to ∼2.2 MDa in two-dimensional Blue Native PAGE (2D BN-PAGE) gels (Huynen et al., 2016). To compare the relative size of TMEM11, we performed 2D BN-PAGE of digitonin-solubilized mitochondria purified from U2OS CRISPRi cells expressing control sgRNA (Fig. 2A). While a small amount of TMEM11 assembled into large complexes consistent with the size of MICOS, the majority of TMEM11 appeared at smaller molecular weights between approximately 60 kDa and 700 kDa.

**Figure 2.**
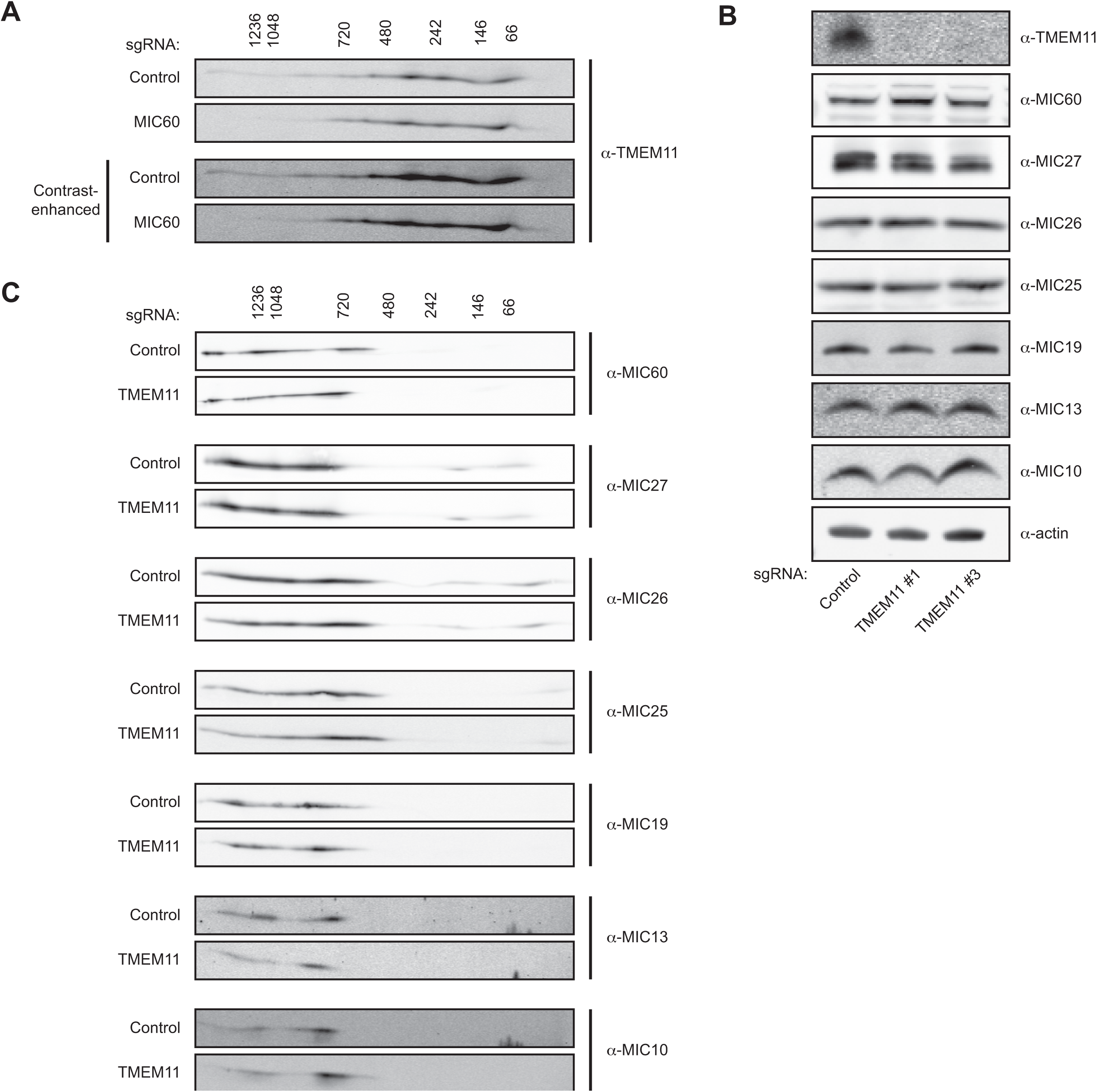
TMEM11 interacts with MICOS but is not required for stability or assembly of the complex. **(A)** Two-dimensional (2D) BN-PAGE and Western analysis of mitochondria isolated from U2OS CRISPRi cells expressing control or TMEM11-targeted sgRNAs and probed with TMEM11 antibody. The molecular weight of assemblies as determined by the first dimension of BN-PAGE are displayed vertically above images. Contrast-enhanced blots are displayed at bottom to enable visualization of higher molecular weight assemblies of TMEM11. **(B)** Western blot analysis of whole cell lysates from cells expressing the indicated sgRNA and probed with the indicated antibodies. **(C)** 2D BN-PAGE and Western analysis from mitochondria isolated from cells expressing the indicated sgRNA as in (A) and probed with the indicated MICOS antibodies.

To determine if the pool of larger TMEM11 assemblies were indeed associated with the MICOS complex, we used multiple sgRNAs to generate cells stably depleted of the core MICOS subunit MIC60 using CRISPRi, loss of which was previously shown to destabilize the MICOS complex (Ott et al., 2015; Stephan et al., 2020). Consistent with published observations, loss of MIC60 led to destabilization of other MICOS subunits and to mitochondrial morphology defects apparent by fluorescence microscopy and EM (Fig. S1). We examined TMEM11 stability in the absence of MIC60 and found that, unlike other MICOS subunits, TMEM11 protein levels were unaffected by MIC60 depletion (Fig. S1). We then purified mitochondria from MIC60-depleted cells (sgRNA #1) and asked whether TMEM11 assembly size was affected in 2D BN-PAGE gels. While the majority of TMEM11 was sized less than ∼700 kDa, there was a reproducible depletion of the minor amount of TMEM11 that migrated at a larger molecular weight (Fig. 2A). These data suggest that a small portion of TMEM11 can stably associate with the assembled MICOS/MIB complex.

Given that a portion of TMEM11 can interact with and assemble into MICOS/MIB complexes, we next asked whether MICOS protein stability was affected in TMEM11-depleted cells (sgRNA #3), which we reasoned could potentially explain the mitochondrial morphology defect observed in the absence of TMEM11. We examined the stability of each MICOS subunit by Western analysis of whole cell lysates obtained from control and TMEM11-depleted cells. However, depletion of TMEM11 did not affect the stability of any MICOS subunit (Fig. 2B).

Given that the majority of TMEM11 assembled into smaller-sized complexes than MICOS, we then considered that these may represent sub-complex assemblies of the MICOS complex and that a role of TMEM11 could be to promote MICOS assembly. In this case, MICOS complex assembly rather than individual subunit stability may be affected in the absence of TMEM11. To test this possibility, we performed 2D BN-PAGE analysis and compared the assembly size of each of the core MICOS subunits in control versus TMEM11-depleted mitochondria. However, we observed no changes in the assembly size of any MICOS subunits (Fig. 2C). Further, in contrast enhanced images, we failed to observe the accumulation of smaller molecular weight complexes in the absence of TMEM11 that may be suggestive of an assembly factor role for TMEM11 (Fig. S2). Altogether, these data indicate that although TMEM11 can assemble into larger molecular weight complexes that require MICOS for their formation, defects in MICOS stability or assembly do not likely explain the mitochondrial morphology defects observed in the absence of TMEM11.

### TMEM11 is an outer mitochondrial membrane protein

To gain insight into the mechanistic role of TMEM11, we dissected its sub-organelle localization and topology at mitochondria. We transduced TMEM11 CRISPRi cells with integrating lentiviral plasmids expressing GFP- or APEX2-GFP-tagged TMEM11 and selected for GFP-expressing cells by FACS. In both cases, TMEM11 was modestly overexpressed compared to endogenous levels (Fig. S3A) and both constructs completely alleviated the mitochondrial morphology defects of TMEM11-depleted cells (Fig. S3B-S3C). We then examined the mitochondrial sub-localization of TMEM11 by performing super-resolution SoRa confocal microscopy of GFP-TMEM11 expressing cells that were fixed and immunolabeled with antibodies targeting either MIC60 or the OMM marker TOMM20, as well as the matrix-localized protein HSP60. While MIC60 was concentrated at discrete focal structures consistent with its known enrichment at cristae junctions (Jans et al., 2013; Stoldt et al., 2019), TMEM11 appeared more uniformly distributed along the membrane (Fig. 3A). However, TMEM11 also occasionally localized to discrete focal structures as compared to both HSP60 and TOMM20 that did not appear to co-localize with MIC60, suggesting it plays a functional role independent of the MICOS complex (Fig. 3A-3B).

**Figure 3.**
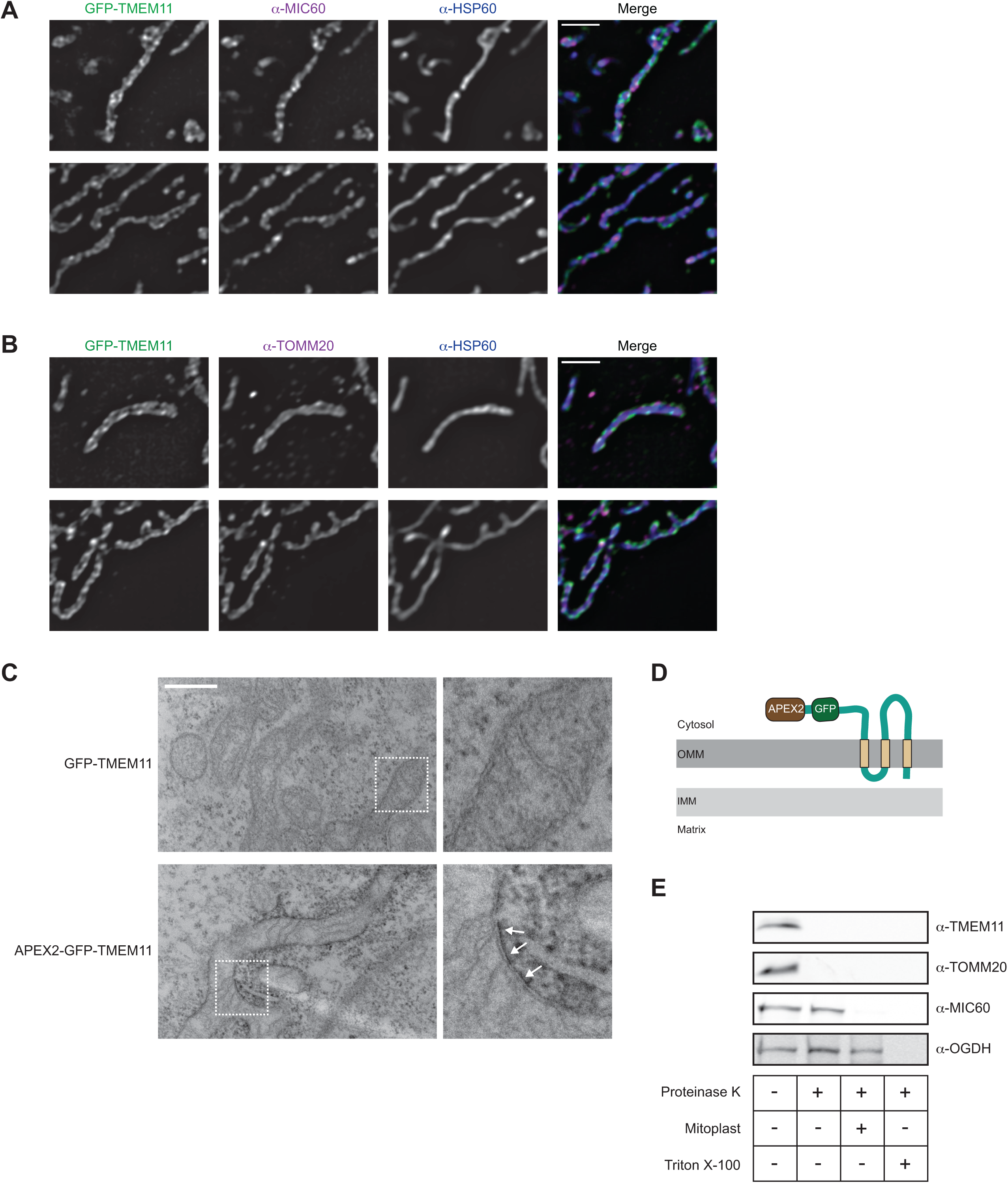
TMEM11 is an outer mitochondria membrane protein. **(A)** Single planes of deconvolved SoRa spinning disk confocal microscopy images are shown of TMEM11 CRISPRi cells expressing GFP-TMEM11 (green) (see Fig. S3) that were fixed and immunolabeled with MIC60 (magenta) and the mitochondrial matrix marker HSP60 (blue). Scale bars = 2 µm. **(B)** As in (A) for cells stained with the outer mitochondrial membrane marker TOMM20 (magenta) and HSP60 (blue). **(C)** Representative electron micrographs are shown from proximity labeling analysis of TMEM11 CRISPRi cells expressing GFP-TMEM11 (top) or APEX2-GFP-TMEM11 (bottom) and treated with DAB and H_2_O_2_ post-fixation. White arrows mark sites of DAB precipitation in cells expressing APEX2-GFP-TMEM11. Enlargements (right) correspond to dotted boxes (left). Scale bar = 500 nm. See also Figure S3D. **(D)** Predicted topology of TMEM11 based on APEX2 proximity labeling (C) and transmembrane domain prediction software. **(E)** Protease protection analysis of mitochondria isolated from wild type U2OS cells. Mitochondria were treated as indicated and Western analysis was performed with the indicated antibodies.

Next, to examine TMEM11 localization relative to mitochondria ultrastructure, we utilized proximity-based APEX labeling to visualize TMEM11 localization on EM images (Lam et al., 2015; Martell et al., 2012). We treated both GFP-TMEM11 and APEX2-GFP-TMEM11 expressing cells post-fixation with 3,3’-diaminobenzidine (DAB) and H_2_O_2_ before subsequent sample preparation for EM. In contrast to GFP-TMEM11 expressing cells, which had no apparent staining, cells expressing APEX2-GFP-TMEM11 formed a dark precipitate near mitochondria in EM sections (Fig. 3C, Fig. S3D). Surprisingly, the DAB precipitate appeared on the exterior of mitochondria, suggesting the APEX2 tag was exposed to the cytosol and consistent with the localization of an OMM protein (Fig. 3D).

As TMEM11 has previously been reported to localize to the IMM, we next sought to re-assess the localization of endogenous TMEM11 using a protease protection assay. Intact mitochondria were isolated from U2OS cells by differential centrifugation and treated with proteinase K before or after osmotic rupture of the OMM. While the IMM protein MIC60 was protected from digestion unless the OMM was ruptured, both TMEM11 and TOMM20 were completely digested by addition of proteinase K to intact mitochondria (Fig. 3E). Altogether, these data indicate that TMEM11 is an OMM protein with a distinct localization from MICOS/MIB complexes.

### TMEM11 forms a complex with BNIP3 and BNIP3L on the mitochondria outer membrane

Given the localization of TMEM11 to the OMM, we next sought to identify TMEM11 interaction partners to understand its functional role. We performed immunoprecipitations and mass spectrometry-based proteomic analysis of lysate from cells stably expressing GFP-TMEM11 with either anti-GFP antibody coupled to Protein G beads or beads alone. Proteins that were identified in control samples were background-subtracted and unique interacting proteins were assigned a normalized spectral abundance factor (NSAF) score accounting for protein molecular weight (Zybailov et al., 2006). Consistent with prior proteomic analysis of MICOS subunits (Guarani et al., 2015) and our 2D BN-PAGE analysis, proteomic analysis of TMEM11 robustly identified multiple subunits of the MICOS complex, including MIC60 and MIC19 (Fig. 4A). However, our analysis also identified several proteins annotated to localize to the OMM. The top scoring interactor was BCL2-Interacting Protein 3-Like (BNIP3L; also known as NIX), a protein implicated in receptor-mediated mitophagy. Other abundant interactors included BNIP3, a BNIP3L paralog, as well as the voltage dependent anion channel (VDAC) family members VDAC1 and VDAC2. BNIP3 and BNIP3L were of particular interest as these proteins were previously identified as reciprocal TMEM11 interactors in large scale yeast two hybrid screens (Luck et al., 2020).

**Figure 4.**
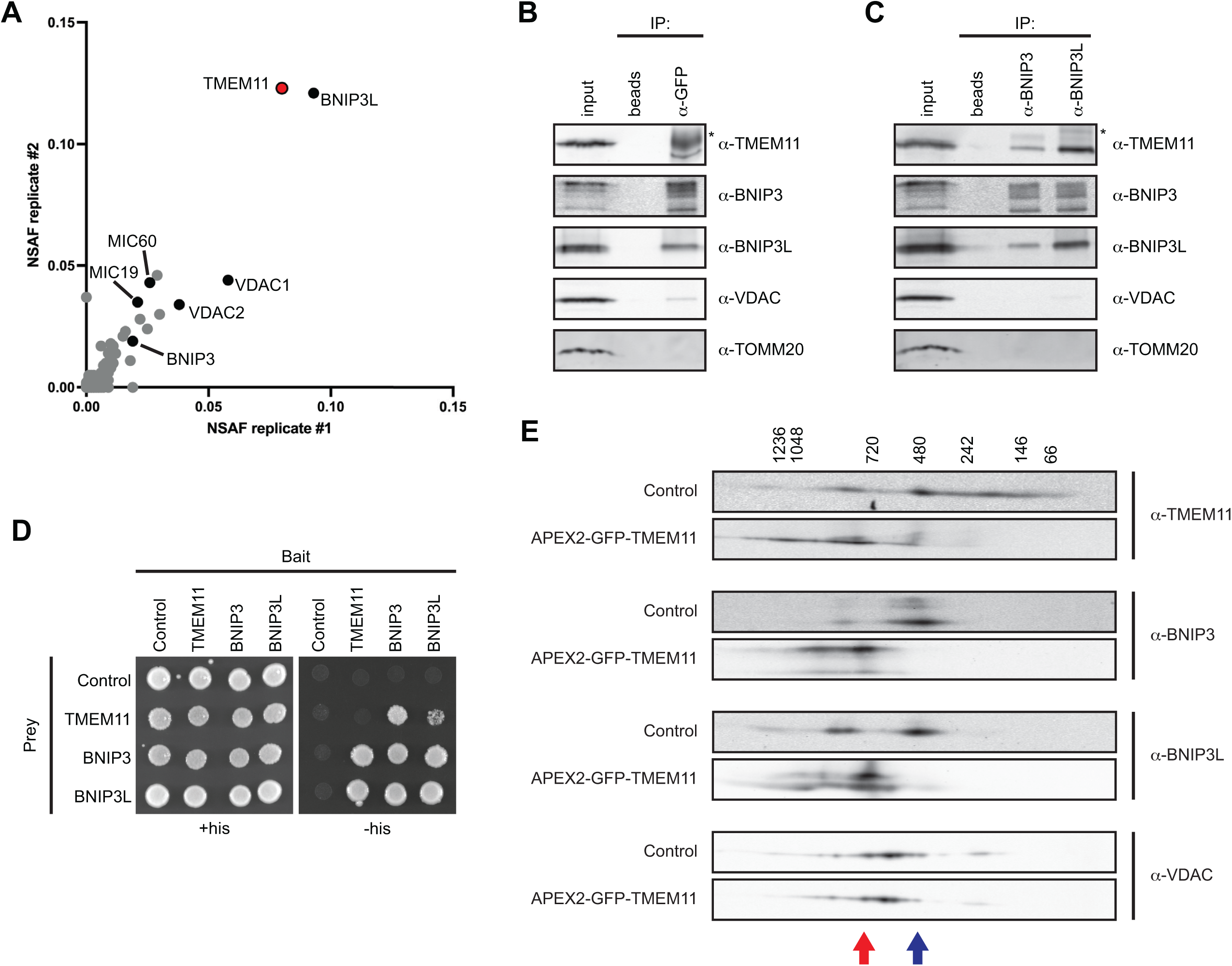
TMEM11 forms a complex with BNIP3 and BNIP3L on the mitochondria outer membrane. **(A)** A plot of normalized spectral abundance factor (NSAF) scores from independent replicates of anti-GFP immunoprecipitation (IP) and mass spectrometry analysis of lysate from TMEM11 CRISPRi cells expressing GFP-TMEM11. **(B)** Western analysis with the indicated antibodies of IP of lysates from GFP-TMEM11-expressing cells with anti-GFP antibody or beads alone. 4% of the total input and 10% of the eluate from each IP were loaded. The asterisk indicates IgG heavy chain. **(C)** IPs were performed as in (B) with anti-BNIP3 and anti-BNIP3L antibodies. **(D)** Yeast two-hybrid analysis of strains expressing the indicated bait and prey proteins and plated on permissive (+his) or selective (-his) media. **(E)** 2D BN-PAGE and Western analysis with the indicated antibodies from mitochondria isolated from U2OS CRISPRi control cells or cells expressing APEX2-GFP-TMEM11, where indicated. Arrows correspond to the position of the peak of TMEM11, BNIP3, and BNIP3L intensity in control (blue) versus APEX2-GFP-TMEM11 cells (red).

To validate the results of our proteomic analysis, we performed immunoprecipitations of lysates from GFP-TMEM11 expressing cells with anti-GFP antibody (Fig. 4B) and immunoblotted for BNIP3L, BNIP3, VDAC1, or TOMM20. TMEM11 robustly interacted with both BNIP3L and BNIP3. While an interaction between TMEM11 and VDAC1 could be detected, this interaction was less robust. TMEM11 also failed to interact with the abundant OMM protein TOMM20, indicating that the interactions between TMEM11 and BNIP3 and BNIP3L were specific. TMEM11 could also be reciprocally identified from lysates from GFP-TMEM11 expressing cells that were immunoprecipitated with BNIP3 or BNIP3L antibodies (Fig. 4C). To determine if TMEM11, BNIP3, and BNIP3L could directly interact, we recapitulated published interactome data (Luck et al., 2020) by expressing each construct in a yeast two-hybrid system. While this assay did not detect TMEM11 self-interaction, BNIP3 and BNIP3L interacted with each other and with TMEM11 (Fig. 4D). Thus, TMEM11 is able to directly interact with BNIP3 and BNIP3L.

We next ascertained whether BNIP3 and BNIP3L were part of similarly-sized molecular weight complexes that we observed for TMEM11 (Fig. 2A). We performed 2D BN-PAGE analysis of mitochondria isolated from U2OS CRISPRi cells expressing control sgRNA and probed for BNIP3 and BNIP3L. Both BNIP3 and BNIP3L appeared most enriched at ∼500 kDa, though migrated as part of both larger and smaller species (Fig. 4E, see blue arrow). While TMEM11 assembled in a wider range of sizes, it also was discretely enriched at as similar size to BNIP3 and BNIP3L. By comparison, VDAC1 predominantly assembled into molecular weight complexes of around ∼600 kDa (Fig. 4E), consistent with previous analysis (Konig et al., 2021).

To further dissect the relationship between TMEM11 and BNIP3/BNIP3L, we took advantage of the increased size of the tandem APEX2-GFP fusion to TMEM11. We reasoned that if BNIP3 and BNIP3L are in a bona fide complex with TMEM11, that their size would correspondingly increase to the increased size of the TMEM11 fusion. Indeed, TMEM11 shifted to a larger molecular weight and was noticeably absent at smaller sizes in 2D BN-PAGE analysis performed on mitochondria isolated from APEX2-GFP-TMEM11 expressing cells (see Fig. 4E, see red arrow). Likewise, both BNIP3 and BNIP3L correspondingly increased in size to a similar sized complex as TMEM11 (Fig. 4E). Importantly, VDAC1, while interacting with TMEM11 in our proteomic analysis, was unaffected by expression of APEX2-GFP-TMEM11 (Fig. 4E). Together, our analyses indicate that TMEM11, BNIP3, and BNIP3L directly interact in an OMM-localized protein complex.

### BNIP3/BNIP3L knockdown alleviates the mitochondrial morphology defects of TMEM11-depleted cells

Given that BNIP3 and BNIP3L are in a complex with TMEM11, we next asked whether the activity of these proteins could be responsible for the altered mitochondrial morphology observed in TMEM11-depleted cells. We performed knockdown of BNIP3 and BNIP3L either individually or in combination by transient siRNA transfection in U2OS CRISPRi cells expressing control sgRNA or TMEM11-targeted sgRNA. We then assayed mitochondrial morphology by staining with Mitotracker and imaging cells by fluorescence microscopy. In control sgRNA cells depleted for BNIP3 or BNIP3L, mitochondria appearance was largely unaffected, although mitochondria tended to be more elongated in BNIP3/BNIP3L double knockdown cells (Fig. 5). Interestingly, examples of mildly enlarged mitochondria (∼10% of control cells) appeared to be significantly and reproducibly alleviated by depletion of BNIP3, but not BNIP3L, indicative of a functionally active role for BNIP3 (Fig. 5A-5B). As before, TMEM11-depleted cells treated with control siRNA had aberrant mitochondria in over half of cells. Remarkably, depletion of BNIP3, and to a lesser extent, BNIP3L, alleviated the mitochondrial morphology defects of TMEM11-depleted cells (Fig. 5A-5B). Combined depletion of BNIP3 and BNIP3L did not additively improve mitochondrial morphology, suggesting other factors may contribute to the defects in mitochondrial morphology observed in the absence of TMEM11. Alternatively, low levels of BNIP3 and/or BNIP3L remaining after siRNA treatment may contribute to the mitochondrial morphology phenotype. Regardless, these data indicate that BNIP3 and BNIP3L activity contributes to mitochondria morphology differences observed in the absence of their binding partner TMEM11.

**Figure 5.**
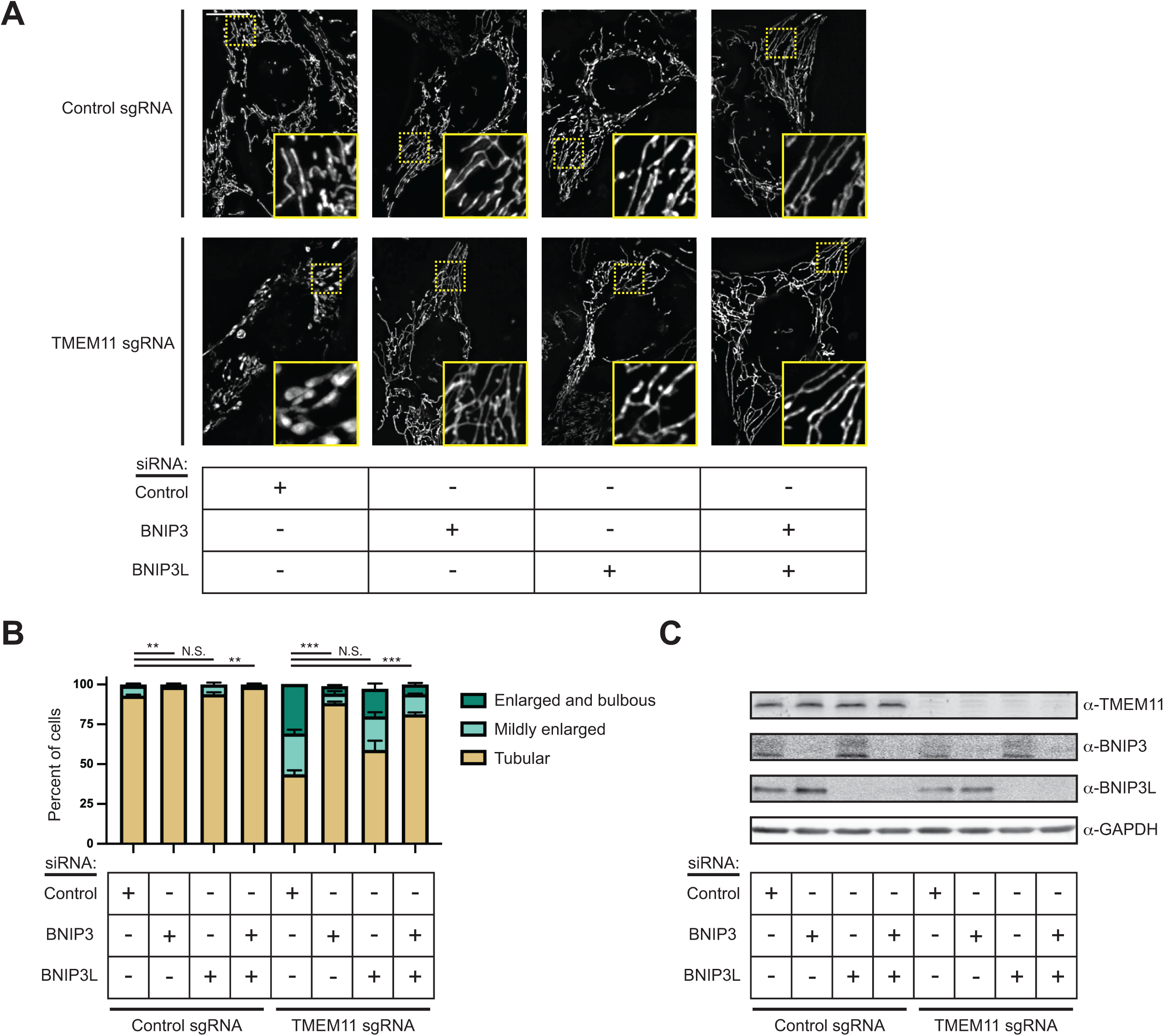
BNIP3/BNIP3L knockdown alleviates the mitochondrial morphology defects of TMEM11-depleted cells. **(A)** Maximum intensity projection fluorescence microscopy images are shown of U2OS CRISPRi cells stably expressing control sgRNA (top) or TMEM11 sgRNA (bottom) that were transiently transfected with the indicated siRNA and stained with Mitotracker Deep Red. Insets correspond to dotted boxes. Scale bar = 15 µm. **(B)** A graph of the categorization of mitochondrial morphology from cells as in (A). Data shown represent 100 cells per condition in each of three independent experiments and bars indicate S.E.M. Asterisks (***p<0.001, **p<0.01) represent unpaired two-tailed *t* test. N.S. indicates not statistically significant. **(C)** Western analysis with the indicated antibodies of whole cell lysates from cells as in (A).

### TMEM11 negatively regulates BNIP3/BNIP3L-mediated basal mitophagy

The primary function of BNIP3 and BNIP3L is thought to be in the turnover of mitochondria through recruitment of LC3 (Liu et al., 2014). BNIP3 and BNIP3L are transcriptionally upregulated to promote their activation, for example, during hypoxia. However, BNIP3 and BNIP3L are expressed at lower levels at steady-state and our data suggests they can contribute to altered mitochondrial morphology when TMEM11 is depleted. Therefore, we asked whether TMEM11-depleted cells are subjected to increased levels of BNIP3- and BNIP3L-mediated mitophagy. We utilized HeLa cells stably expressing the mitophagy reporter mito-mKeima (Lazarou et al., 2015). mKeima is a pH-sensitive fluorophore that, when targeted to the mitochondrial matrix, enables the differential visualization of both active mitochondria as well as those that are targeted to lysosome for degradation depending on the excitation wavelength (Sun et al., 2017). To determine the contribution of TMEM11 to steady-state basal mitophagy, we imaged HeLa mito-mKeima cells transfected with control siRNA or siRNA targeting TMEM11. Images were then manually scored for the number of acidified mitochondrial puncta that could be visualized per cell, an indicator that the mitochondria were targeted to lysosomes. Even in the absence of mitophagy stimuli, approximately 50% of cells had at least one acidic mitochondrial puncta, suggesting that HeLa mito-mKeima cells undergo basal mitophagy (Fig. 6A-6C). Consistent with our results in U2OS CRISPRi cells, depletion of TMEM11 by siRNA in HeLa mito-mKeima cells caused mitochondria to become more enlarged and bulbous in appearance (Fig. 6A, 6D). Strikingly, TMEM11-depleted cells had significantly higher numbers of acidified mitochondria than in control cells (12% control vs. 22% of TMEM11-depleted cells had 10 or more puncta; Fig. 6B-6C). We co-transfected both control and TMEM11 siRNA-treated cells with siRNAs targeting BNIP3 and/or BNIP3L to determine if the increased prevalence of mitophagy in TMEM11-depleted cells was related to their function. Remarkably, acidified mitochondria puncta were greatly reduced in the absence of BNIP3 or BNIP3L and nearly abolished in the absence of both proteins in either control or TMEM11-depleted cells (Fig. 6A-6C). These data indicate that BNIP3 and BNIP3L contribute to low levels of steady-state mitophagy in HeLa mito-mKeima cells. Further, depletion of the BNIP3/BNIP3L interactor TMEM11 leads to an increase in BNIP3/BNIP3L-dependent mitophagy.

**Figure 6.**
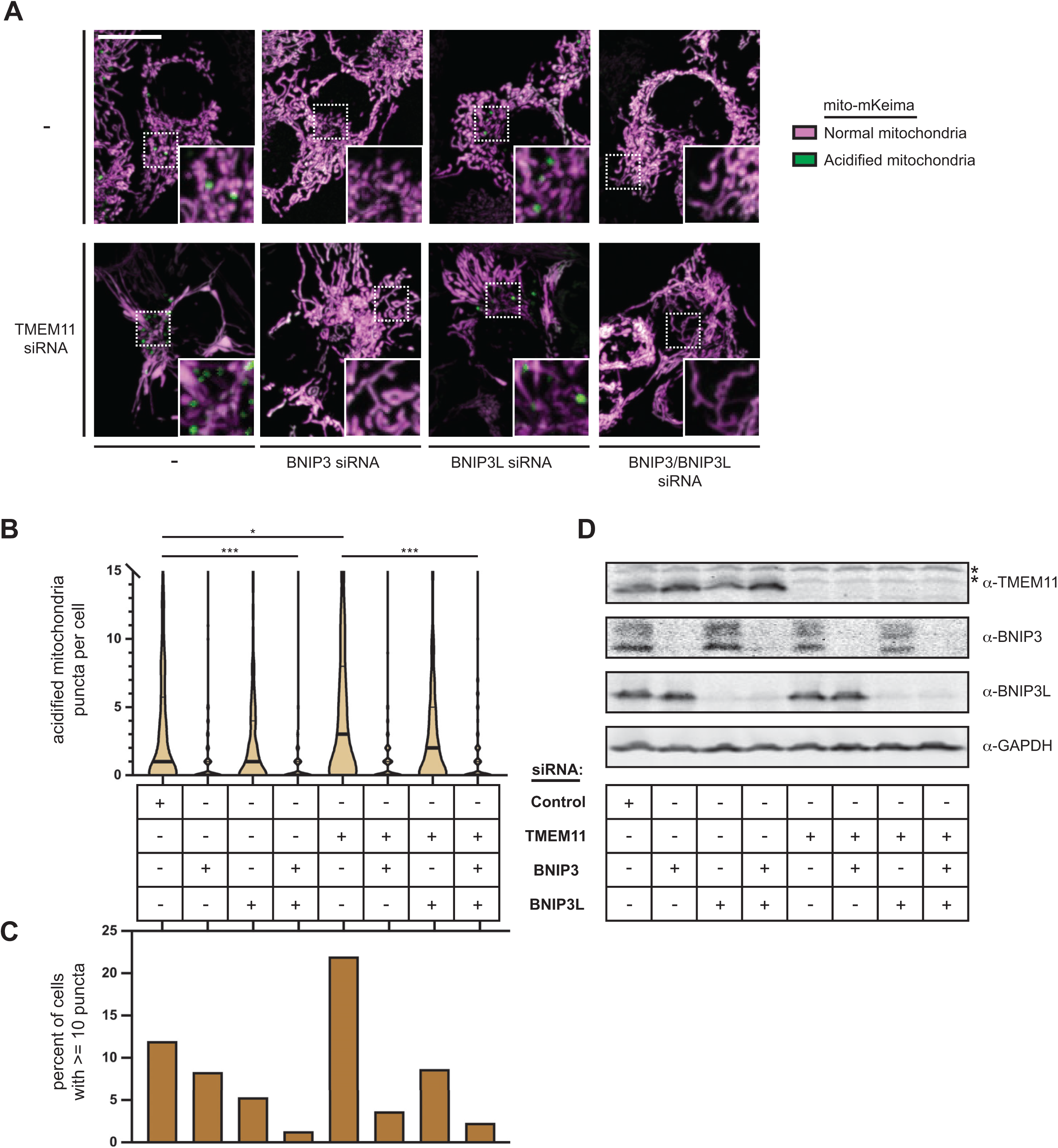
TMEM11 negatively regulates BNIP3/BNIP3L-mediated basal mitophagy. **(A)** Merged maximum intensity projections of confocal fluorescence microscopy images of HeLa mito-mKeima expressing cells that were transiently transfected with siRNAs targeting TMEM11 (bottom row) and BNIP3 and/or BNIP3L, where indicated, and excited with a 471 nm laser (magenta, neutral pH mitochondria) and a 561 nm laser (green, acidified mitochondria). Scrambled control siRNA was used in cases with no other target. Insets correspond to dotted boxes. Scale bars = 15 µm. **(B)** A violin plot depicting the number of acidified mitochondria puncta per cell corresponding to green labeling from cells with the indicated siRNA treatments as in (A). Data shown represent the summation of three independent experiments with 100 cells from each experiment. Asterisks (***p<0.001, *p=0.013) represent unpaired two-tailed *t* test. Bold horizontal lines mark medians and thin horizontal lines mark quartiles for each condition. For clarity, the small number of cells with more than 15 puncta are not depicted. **(C)** A histogram of the percent of cells from each condition as in (A, B) with at least 10 puncta. **(D)** Western analysis with the indicated antibodies of whole cell lysates from cells as in (A) treated with the indicated siRNAs. The asterisks indicate cross-reacting bands.

### TMEM11 depletion sensitizes cells to BNIP3-mediated mitophagy during hypoxia

Next, to examine the contribution of TMEM11 to the regulation of BNIP3 and BNIP3L during HIF1-*α*-stimulated mitophagy, we exposed cells to the hypoxia mimetic CoCl_2_ (250µM for 24h). As expected (Bellot et al., 2009; Sulkshane et al., 2021), BNIP3 and BNIP3L protein stability increased upon CoCl_2_ treatment in HeLa mito-mKeima expressing cells. In contrast, TMEM11 stability was unaffected by treatment (Fig. 7A).

**Figure 7.**
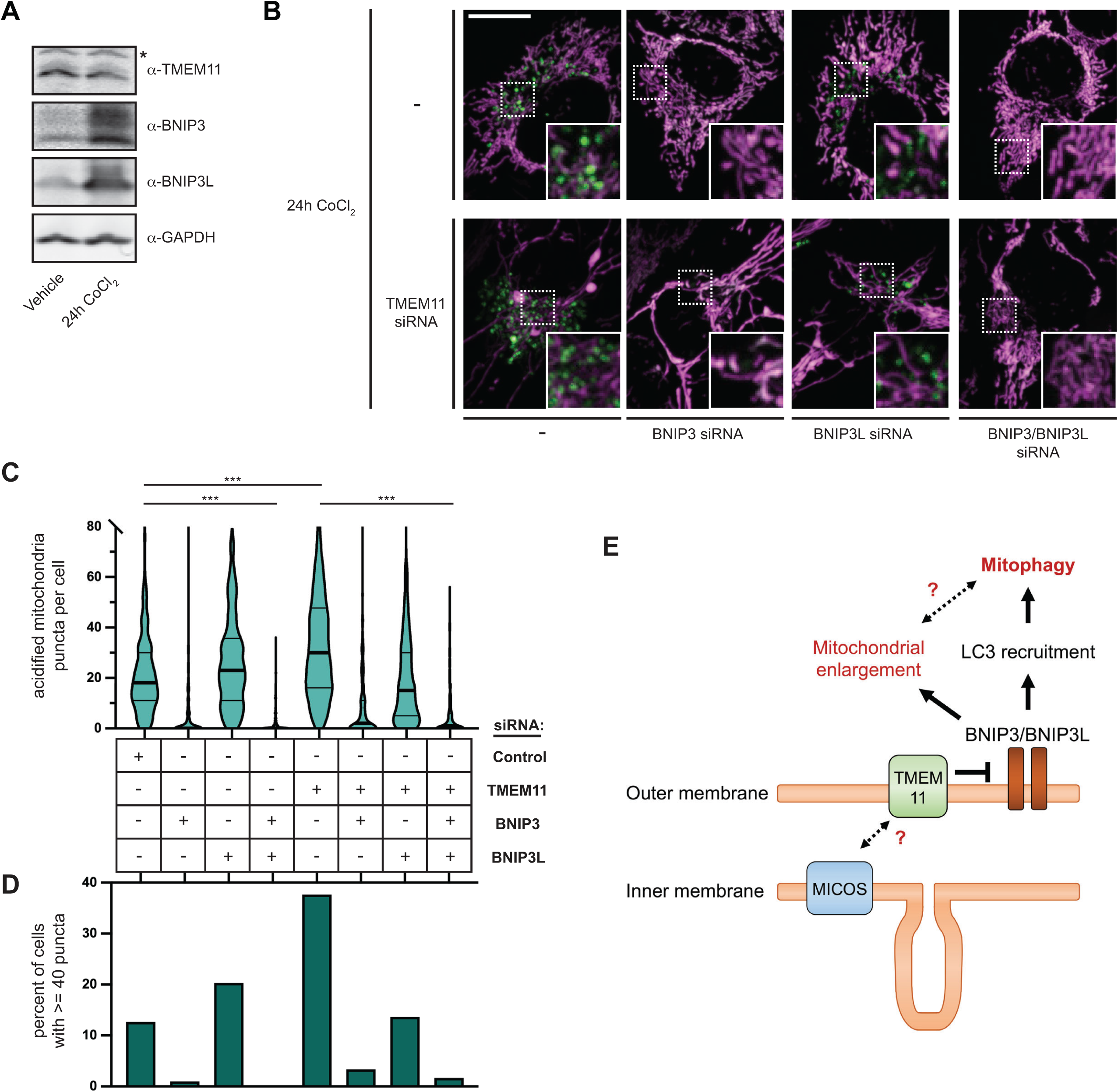
TMEM11 depletion sensitizes cells to BNIP3-mediated mitophagy during hypoxia. **(A)** Western blot analysis with the indicated antibodies of whole cell lysates from Hela mito-mKeima expressing cells treated with vehicle or CoCl_2_ (250 µM, 24h). The asterisk indicates a cross-reacting band. **(B)** Merged maximum intensity projections of confocal fluorescence microscopy images of HeLa mito-mKeima expressing cells that were transiently transfected with siRNAs targeting TMEM11, BNIP3, and/or BNIP3L, where indicated, treated with CoCl_2_ (250 µM, 24h), and excited with a 471 nm laser (magenta, neutral pH mitochondria) and a 561 nm laser (green, acidified mitochondria). BNIP3 and/or BNIP3L-silenced cells were simultaneously treated with Q-VD-OPh to prevent apoptosis. Scrambled control siRNA was used in cases with no other target. Insets correspond to dotted boxes. Scale bars = 15 µm. **(C)** A violin plot depicting the number of acidified mitochondria puncta per cell corresponding to green labeling from cells with the indicated siRNA treatments as in (B). Data shown represent the summation of three independent experiments with 100 cells from each experiment. Asterisks (***p<0.001) represent unpaired two-tailed *t* test. Bold horizontal lines mark medians and thin horizontal lines mark quartiles for each condition. For clarity, the small percentage of cells with more than 80 puncta are not depicted. **(D)** A histogram of the percent of cells from each condition as in (B, C) with at least 40 acidified mitochondrial puncta. **(E)** A model for the role of TMEM11 as a negative regulator of BNIP3/BNIP3L-mediated mitochondrial enlargement and mitophagy.

Next, we examined mitophagy during CoCl_2_ treatment in HeLa cells stably expressing mito-mKeima and transfected with scrambled control siRNA. Consistent with the increase in BNIP3 and BNIP3L protein stability, CoCl_2_ treatment stimulated mitophagy and drastically increased the number of acidified mito-mKeima labeled mitochondria in each cell (the median number of acidified puncta increased from 1 to 18 after treatment; compare Fig. 6B and 7C). To determine whether the increase in mitophagy was dependent on BNIP3 and/or BNIP3L, we performed knockdowns with siRNA and co-treated with the caspase inhibitor Q-VD-OPh to prevent apoptotic cell death (Bellot et al., 2009). BNIP3 was largely required for CoCl_2_-induced mitophagy, though combined depletion of both BNIP3 and BNIP3L nearly completely abrogated mitochondrial acidification post-treatment (Fig. 7B-7D).

Given that TMEM11 negatively regulated BNIP3/BNIP3L-dependent mitophagy at steady-state and that BNIP3/BNIP3L are transcriptionally upregulated to drive mitophagy upon CoCl_2_ treatment while TMEM11 remains stable, we next asked whether depletion of TMEM11 would further sensitize mitochondria to mitophagy upon treatment. Remarkably, TMEM11-depleted cells had drastically higher numbers of acidified mito-mKeima labeled mitochondria compared to control cells 24h after CoCl_2_ treatment (Fig. 7B-7D). In particular, cells with abundant mitophagosomes could be more readily observed in the absence of TMEM11 (13% of control cells vs. 38% of TMEM11-depleted cells had at least forty acidified mitochondrial puncta; Fig. 7D). The increased mitophagy in the absence of TMEM11 was also primarily alleviated by depleting cells of BNIP3, however, was further reduced by silencing of both BNIP3 and BNIP3L (Fig. 7B-7D). Thus, depletion of the BNIP3/BNIP3L-interacting partner TMEM11 sensitized cells to BNIP3/BNIP3L-driven mitophagy under hypoxia-mimicking conditions.

## Discussion

Although PINK1/Parkin-mediated mitophagy is better mechanistically understood, mitochondrial turnover by BNIP3/BNIP3L-dependent receptor-mediated mitophagy is a critical modulator of mitochondrial turnover in the heart, under stress conditions such as hypoxia, during development, and is frequently mis-regulated in cancer cells (Macleod, 2020; Ney, 2015). However, a long-standing question has been how BNIP3/BNIP3L activity is regulated at steady-state. Our data reveal that the mitochondrial morphology regulator TMEM11, while originally thought to be a mitochondrial IMM auxiliary component of the MICOS complex, localizes to the OMM where it forms a complex with BNIP3 and BNIP3L. Further, we find that TMEM11 negatively regulates BNIP3/BNIP3L-dependent receptor-mediated mitophagy both at basal and under hypoxia-mimetic conditions. These data are consistent with the model that the TMEM11/BNIP3/BNIP3L complex regulates the homeostasis of receptor-mediated mitophagy (Fig. 7E).

While our data suggest that BNIP3 and BNIP3L both directly interact with TMEM11 and associate in higher molecular-weight complexes together, mitochondrial morphology and hypoxia defects that occur in the absence of TMEM11 were primarily alleviated by depleting BNIP3 and not BNIP3L. Further, depletion of TMEM11 increased mitophagy, though this increase was primarily due to BNIP3 activity. Thus, while our findings are consistent with the model that TMEM11 negatively regulates BNIP3/BNIP3L activity, it is clear additional factors must contribute to positively modulating their function. Indeed, BNIP3L is upregulated during CoCl_2_ treatment, but this alone is insufficient to lead to a drastic increase in mitophagy in the absence of BNIP3. Both BNIP3 and BNIP3L are extensively post-translationally modified and these modifications have been shown to be necessary to regulate mitophagy (Poole et al., 2021; Rogov et al., 2017; Zhu et al., 2013). Additional interacting proteins may also play a role. One possibility is that distinct types of stress lead to differential activation of BNIP3 versus BNIP3L-dependent basal mitophagy. Meanwhile, BNIP3L is specifically upregulated to eliminate mitochondria from maturing erythrocytes and keratinocytes (Sandoval et al., 2008; Schweers et al., 2007; Simpson et al., 2021), and an outstanding question is whether TMEM11 plays an inhibitory role in these processes.

Our data also indicate that BNIP3 activity leads to a mitochondrial morphology defect that occurs in the absence of TMEM11. Interestingly, BNIP3 and BNIP3L have been previously associated with hypoxia-associated mitochondria enlargement, at least in certain cell types (Chiche et al., 2010). Stimulation of mitophagy by CoCl_2_ treatment in HeLa mito-Keima cells also empirically causes an apparent increase in mitochondrial enlargement, however only in a minor subset of cells. We also observe that under basal conditions, a small percentage of U2OS cells appear to have mildly enlarged mitochondria that are also reproducibly alleviated by depletion of BNIP3 (Fig. 4B). However, the functional relationship between BNIP3/BNIP3L-dependent mitochondrial enlargement and mitophagy is not clear as the morphology defect in the absence of TMEM11 is not strictly correlated with the amount of mitophagy induction. Chiche et al. propose that the BNIP3/BNIP3L-dependent enlargement of mitochondria they observed during hypoxia protects against apoptotic induction. An additional possibility is the increased diameter of the enlarged mitochondria may protect functional parts of the mitochondrial network against selective mitophagy.

Finally, our data suggest that TMEM11 not only physically interacts with BNIP3 and BNIP3L, but also interacts to a lesser degree in higher molecular weight complexes with the MICOS/MIB complex. TMEM11 does not appear to affect MICOS stability or assembly, begging the question of whether the MICOS-TMEM11 interaction plays a functional role independently of the TMEM11/BNIP3/BNIP3L complex, or whether cristae organization and mitophagy are linked. An interesting possibility that remains to be tested in the future is whether TMEM11 may sense mitochondrial dysfunction through its interaction with MICOS as a way of modulating BNIP3/BNIP3L activity.

## Materials and Methods

### Cell culture

U2OS cells and derivatives, HeLa mito-mKeima cells (a kind gift of Richard Youle; (Lazarou et al., 2015)), and HEK 293T cells were cultured in DMEM (Sigma D5796) supplemented with 10% fetal bovine serum (Sigma), 25 mM HEPES, 100 units/mL penicillin, and 100 µg/mL streptomycin. All experiments using CRISPRi cells were performed on early passages (<10) after sorting. Cell lines were routinely tested for mycoplasma contamination.

### Plasmids and siRNA oligonucleotides

Individual TMEM11 sgRNA and MIC60 sgRNA vectors were generated by annealing the following forward and reverse oligonucleotides and ligating into pU6-sgRNA Ef1α-Puro-T2A-BFP (Horlbeck et al., 2016) digested with BstXI and BlpI.

TMEM11 sgRNA #1 forward: 5’-TTGGGAAGGAGGCGTCTTGGCCCGTTTAAGAGC-3’

TMEM11 sgRNA #1 reverse: 5’-TTAGCTCTTAAACGGGCCAAGACGCCTCCTTCCCAACAAG-3’

TMEM11 sgRNA #3 forward: 5’-TTGGCGAGAGAGGTGAGATCCAAGTTTAAGAGC-3’

TMEM11 sgRNA #3 reverse: 5’-TTAGCTCTTAAACTTGGATCTCACCTCTCTCGCCAACAAG-3’

MIC60 sgRNA #1 forward: 5’-TTGGCGCGGCGGCGCGAGTTAAGGTTTAAGAGC-3’

MIC60 sgRNA #1 reverse: 5’-TTAGCTCTTAAACCTTAACTCGCGCCGCCGCGCCAACAAG-3’

MIC60 sgRNA #2 forward: 5’-TTGGTGGTGGACTCGAGCTGCCGGTTTAAGAGC-3’

MIC60 sgRNA #2 reverse: 5’-TTAGCTCTTAAACCGGCAGCTCGAGTCCACCACCAACAAG-3’

pLVX-Puro GFP-TMEM11 was generated by PCR amplifying TMEM11 from human cDNA and cloning into the XhoI/BamHI sites of pAcGFP1-C1 (Takara), followed by subsequent cloning of the GFP-TMEM11 cassette into the Xho/BamHI sites of pLVX-Puro (Takara) by isothermal assembly. pLVX-Puro APEX2-GFP-TMEM11 was generated by PCR amplifying the GFP-TMEM11 cassette and the APEX2 cassette from pcDNA3 Connexin43-GFP-APEX2 (kindly provided by Alice Ting; Addgene #49385) (Lam et al., 2015) and cloning into the Xho/BamHI sites of pLVX-Puro by isothermal assembly.

pGADT7-TMEM11 and pGBKT7-TMEM11 were generated by cloning the TMEM11 cassette from pGFP-TMEM11 and ligating it into NdeI/BamHI sites of pGADT7 and pGBKT7 plasmids (Takara), respectively, by isothermal assembly. pGADT7-BNIP3 and pGBKT7-BNIP3 were generated by cloning the BNIP3 cassette from pDEST40-BNIP3 (a gift from Angelique Whitehurst) and ligating it into NdeI/BamHI sites in pGADT7 AD and pGBKT7, respectively, by isothermal assembly. pGADT7-BNIP3L and pGBKT7-BNIP3L were generated by cloning the BNIP3L cassette from pDEST40-BNIP3L (a gift from Angelique Whitehurst) and ligating it into NdeI/BamHI sites in pGADT7 AD and pGBKT7, respectively, by isothermal assembly.

For siRNA depletion, Silencer Select siRNAs (Negative control no. 2 – 4390846; TMEM11 - s16855; BNIP3 – s2060; BNIP3L s2063; ThermoFisher) were used for all treatments

### Lentivirus production and generation of stable cell lines

Lentivirus of sgRNA-expressing plasmids and TMEM11-expressing plasmids was generated as previously described (Le Vasseur et al., 2021). Briefly, HEK293Ts were transfected with standard packaging plasmids (0.1 µg of pGag/Pol, 0.1 µg of pREV, 0.1 µg of pTAT, and 0.2 µg of pVSVG) and 1.5 µg of lentiviral vector using TransIT-LT1 Transfection Reagent (Mirus). Lentiviral supernatant was harvested and filtered through a 0.45 µm PES filter.

To generate CRISPRi stable lines, U2OS dCas9 cells were seeded into a six-well plate (200,000 cells per well) and incubated for 24h with 750 µL viral supernatant of sgRNA-expressing plasmids supplemented with 5.33 µg/mL polybrene. After infection, cells were allowed to grow and the top 50% of TagBFP expressing cells were sorted by FACS in the UTSW Flow Cytometry Core Facility. Control sgRNA-expressing U2OS CRISPRi cells were previously described (Le Vasseur et al., 2021).

GFP-TMEM11 and APEX2-GFP-TMEM11 lentiviral plasmids were transduced as above (100µL viral supernatant) into U2OS CRISPRi cells stably expressing TMEM11 sgRNA #3. The bottom 33% of GFP-expressing cells were sorted by FACS in the UTSW Flow Cytometry Core.

### Whole cell lysates and Western analysis

To prepare whole cell lysates, cells were trypsinized, harvested, washed once with DPBS, and lysed in 1x RIPA buffer (150 mM NaCl, 50 mM Tris HCl pH7.5, 1% Na-deoxycholate, 0.1% SDS, 1% NP-40, 1 mM EDTA) supplemented with 1x protease inhibitor cocktail (MilliporeSigma 539131). Protein concentration was determined using a Bradford assay and normalized before adding 6x Laemmli buffer (0.66% SDS, 24% glycerol, 0.2 M Tris HCl pH 6.8, 0.01% bromophenol blue, 10% beta-mercaptoethanol) to a final concentration of 1x. Samples were heated for 5 min at 95°C and equal amounts of proteins were loaded on Tris-Glycine or Tricine polyacrylamide gels based on protein size. After electrophoresis, proteins were electroblotted on PVDF membranes (0.45 µm pore size) or nitrocellulose membranes (0.2 µm pore size), and immunoblotted with the following primary antibodies: anti-TMEM11 (Proteintech 16564-1-AP), anti-actin (Proteintech 66009-1-Ig), anti-MIC10 (abcam 84969), anti-MIC13 (Proteintech 25515-1-AP), anti-MIC19 (Atlas antibodies HPA042935), anti-MIC25 (Proteintech 20639-1-AP), anti-MIC26 (ThermoFisher MA515493), anti-MIC27 (ThermoFisher PA5-51427), anti-MIC60 (Proteintech 10179-1-AP), anti-OGDH (Proteintech 15212-1-AP), anti-BNIP3 (Santa Cruz sc-56167 or Cell Signaling Technology 44060), anti-BNIP3L (Cell Signaling Technology 12396), anti-VDAC1/2 (Proteintech 10866-1-AP), anti-GAPDH (Proteintech 60004-1-Ig), anti-TOMM20 (Abcam 56783). The appropriate secondary antibodies conjugated to DyLight 680 or DyLight 800 (Thermo Fisher Scientific) were used and visualized with the Odyssey Infrared Imaging System (LI-COR). Linear adjustments (and nonlinear adjustments for contrast enhancement where stated) to images were made using Adobe Photoshop.

### Mitochondria isolation

Mitochondria were isolated by differential centrifugation as previously described (Hoppins et al., 2011b) with the following modifications. U2OS cells were grown to confluency on 15 cm dishes, rinsed with DPBS, and harvested by scraping into warm DPBS and centrifugation (200x g, 5 min). Cells were resuspended in 5-10 pellet volumes of cold mitochondria isolation buffer (10 mM Tris/MOPS pH 7.4, 0.25 M Sucrose, 1 mM EGTA) and lysed with 25 strokes of a glass Dounce homogenizer fitted with a tight pestle. Unbroken cells and nuclei were pelleted by centrifugation (600x g, 10 min, 4°C). This process was repeated, and cellular lysate was pooled. Crude mitochondria were then isolated by centrifugation (10000x g, 15 minutes, 4°C) and resuspended in cold mitochondria isolation buffer. Mitochondria concentration was measured by a Bradford assay and 100 µg aliquots were flash frozen in liquid nitrogen and stored at -80°C.

### 2D BN-PAGE analysis

Mitochondria aliquots (100 µg) were thawed on ice, pelleted by centrifugation (21000x g, 10 min, 4°C), and resuspended in 20 µL of 1x NativePAGE Sample Buffer (ThermoFisher) supplemented with 1x protease inhibitor cocktail and digitonin (SigmaMillipore) to a final detergent:protein ratio of 6 g/g. Samples were solubilized on ice for 15 min and subjected to centrifugation (21000x g, 30 min, 4°C). The supernatant containing solubilized mitochondria was supplemented with Coomassie Blue G-250 dye to a final detergent:dye ratio of 16 g/g before running on a 3-12% NativePAGE Mini Protein Gel (ThermoFisher) according to manufacturer’s directions. Protein complex sizes were standardized with NativeMark Unstained Protein Standard (ThermoFisher). For the second dimension SDS-PAGE, entire lanes were excised and incubated in 10 mL of denaturing buffer (0.12 M Tris-HCl pH 6.8, 4% SDS, 20% glycerol and 10% beta-mercaptoethanol) for 25 min (Fiala et al., 2011). Gel slices were microwaved for 10 seconds halfway through incubation. Each gel slice was loaded horizontally on a denaturing Tris-Glycine polyacyrlamide gel and sealed into position by overlaying with 0.75% agarose in SDS-PAGE running buffer. Western blotting was performed as described above.

### Protease protection assay

Mitochondria were isolated from U2OS cells by differential centrifugation as described above, except cells were lysed with a single set of 10 strokes with a glass Dounce homogenizer and crude mitochondria were obtained by lower speed centrifugation (7400x g, 10 min, 4°C). Protease protection analysis was performed as previously described (Hoppins et al., 2011a) with the following modifications. Mitochondria (25 µg) were resuspended in 500µL mitochondria isolation buffer (10 mM Tris/MOPS pH 7.4, 0.25 M Sucrose, 1 mM EGTA), mitoplast/swelling buffer (10 mM Tris/MOPS, pH 7.4), or solubilization buffer (mitochondria isolation buffer containing 1% Triton X-100). The mitoplast/swelling sample was incubated 15 minutes on ice and then vigorously pipetted 15 times to disrupt the OMM. Proteinase K (100 µg/mL) was then added to the indicated samples and incubated on ice for 15 min. Protease digestion was stopped by addition of PMSF (2 mM) and incubating the samples on ice for 5 min. The Triton X-100 solubilized sample was immediately subjected to TCA precipitation (12.5%). The remaining samples were subjected to centrifugation (10400x g, 15 min, 4°C), supernatants were discarded, and pellets were resuspended in 50 µL mitochondria isolation buffer supplemented with 1x protease inhibitor cocktail. Proteases were denatured (65°C, 10 min) and samples were TCA precipitated. All protein pellets were washed in acetone, dried, and resuspended in 1x MURB sample buffer (100 mM MES pH 7.0, 3 M urea, 1% SDS, 10% beta-mercaptoethanol) prior to Western analysis.

### Immunoprecipitations

U2OS TMEM11 (sgRNA #3) CRISPRi cells expressing GFP-TMEM11 were grown to confluency in 15 cm dishes and harvested by trypsinization and centrifugation. Cells were washed in DPBS and lysed by incubation for 30 min on ice in three pellet volumes of immunoprecipitation lysis buffer (IPLB) (20 mM HEPES-KOH pH7.4, 150 mM KOAc, 2 mM Mg(Ac)2, 1 mM EGTA, 0.6 M sorbitol) supplemented with 1% digitonin and 1x protease inhibitor cocktail. Cell lysate was then recovered after centrifugation (11500x g, 10 min, 4°C) and protein concentration was measured with a Bradford assay before lysate was stored at -80°C.

For mass spectrometry, immunoprecipitation was performed on two independently prepared samples. Equivalent amounts of thawed lysate (5mg replicate 1, 10mg replicate 2) were incubated for 4 h at 4°C with 5 µg anti-GFP antibody (Abcam ab290) or mock-treated as a control. Antibodies were captured with 100 µL of μMACS protein G beads (Miltenyi) for 4h at 4°C. Beads were isolated with μ columns and a μMACS separator (Miltenyi), washed three times with 800 µL of IPLB supplemented with 0.1% w/v digitonin and 1x protease inhibitor cocktail, and two times with 500 µL of IPLB. Samples were eluted using on-bead trypsin digestion with 25 µL of elution buffer-I (2 M urea, 50 mM Tris-HCl pH 7.5, 1 mM DTT, 5 µg/mL trypsin) for 30 minutes followed by 2x 50 µL elution buffer-II (2 M urea, 50 mM Tris-HCl pH 7.5, 5 mM 2-chloroacetamide) and incubated overnight. Samples were quenched by addition of 1 µL trifluoroacetic acid and submitted to the UT Southwestern Proteomics Core for liquid chromatography/tandem MS analysis. Analysis of the samples and raw MS data files were performed as previously described (Tirrell et al., 2020) except peptide identification was performed against the *Homo sapiens* protein database from UniProt. Abundance values for each identified protein were calculated as the sum of the peak intensities for each peptide identified for that protein. Proteins were included in analysis that were identified with >40-fold abundance enrichment in GFP-treated samples relative to beads alone and had at least 5 peptide spectral matches in either experiment. Normalized spectral abundance factor (NSAF) was then calculated for each replicate (the total number of spectral counts (SpC) identifying a protein, divided by the protein’s length (*L*), divided by the sum of SpC/*L* for all proteins in each experiment (Zhu et al., 2010).

For Western analysis, immunoprecipitations were performed as above using 1.4 mg lysate per each sample. The following amounts of antibody were used: 2.5 µg anti-GFP (Abcam ab290), 0.07 µg anti-BNIP3 (Cell Signaling Technology 44060), and 0.92 µg anti-BNIP3L (Cell Signaling Technology 12396). Antibodies were captured with 25 µL of μMACS protein G beads. Proteins were eluted with 2x 25 µL of 2x Laemmli buffer pre-warmed to 95°C.

### Yeast two hybrid analysis

Yeast two-hybrid analysis was performed with the Matchmaker Gold Yeast Two-Hybrid System (Takara). Y2H Gold and Y187 yeast strains were transformed with bait (pGBKT7 plasmids and derivatives) and prey (pGAD T7 plasmids and derivatives), respectively, by lithium acetate transformation. Haploid bait- and prey-expressing strains were mated on YPD plates (1% yeast extract, 2% peptone, 2% glucose) for 24 hours and diploids were subsequently selected on synthetic dextrose (SD; 0.7% yeast nitrogen base, 2% glucose, amino acids) -leu-trp plates. Cells were grown to exponential phase in SD-leu-trp media, normalized to 0.5 OD600 per mL, and cells were spotted on SD-leu -trp (permissive) and SD-leu-trp-his (selection) plates. Plates were then incubated at 30°C prior to analysis.

### Immunofluorescence analysis of GFP-TMEM11 expressing cells

Cells were grown to ∼50% confluency in glass bottom dishes (Cellvis). Cells were fixed in 4% paraformaldehyde solution in PBS (15 minutes, room temperature). Fixed cells were permeabilized (0.1% Triton X-100 in PBS), blocked (10% FBS and 0.1% Triton X-100 in PBS), and then incubated with the indicated primary antibodies (anti-TOMM20 (Abcam 56783), anti-HSP60 (Proteintech 15282-1-AP), and/or anti-MIC60 (Abcam, cat# 110329)) and secondary antibodies (anti-rabbit Alexa 647 Plus (ThermoFisher PIA32795) and donkey anti-mouse Alexa 555 (ThermoFisher A-31570)) in blocking buffer. The subcellular localization of proteins was visualized on a Nikon Ti2 microscope equipped with Yokogawa CSU-W1 spinning disk confocal and SoRa modules, a Hamamatsu Orca-Fusion sCMOS camera and a Nikon 100x 1.45 NA objective. Z-series images were acquired using the SoRa module (additional 2.8 optical magnification) using a 0.2 µm step size. Images were deconvolved using AutoQuant X3 (10 iterations, blind deconvolution, and low noise), and linear adjustments were made with Fiji.

### Electron microscopy analysis

To determine mitochondrial ultrastructure in CRISPRi cells, 100,000 cells were plated onto glass-bottom dishes (MatTek), allowed to adhere for ∼16h, and fixed with 2.5% (v/v) glutaraldehyde in 0.1M sodium cacodylate buffer and submitted to UTSW Electron Microscopy Core Facility for further processing. After five rinses in 0.1 M sodium cacodylate buffer, they were post-fixed in 1% osmium tetroxide and 0.8 % K_3_[Fe(CN_6_)] in 0.1 M sodium cacodylate buffer for 1h at 4°C. Cells were rinsed with water and en bloc stained with 2% aqueous uranyl acetate overnight at 4°C. After five rinses with water, specimens were dehydrated with increasing concentration of ethanol at 4°C, infiltrated with Embed-812 resin and polymerized in a 60°C oven overnight. Embed-812 discs were removed from MatTek plastic housing by submerging the dish in liquid nitrogen. Pieces of the disc were glued to blanks with super glue and blocks were sectioned with a diamond knife (Diatome) on a Leica Ultracut UCT (7) ultramicrotome (Leica Microsystems) and collected onto copper grids and post-stained with 2% uranyl acetate in water and lead citrate. Images were acquired on a JEM-1400 Plus transmission electron microscope equipped with a LaB_6_ source operated at 120 kV using an AMT-BioSprint 16M CCD camera.

For proximity labeling, cells stably expressing APEX2-GFP-TMEM11 or GFP-TMEM11 were processed as previously described (Datta et al., 2019) with the following modifications. Briefly, 50,000 cells were plated on gridded glass bottom dishes (MatTek) and allowed to adhere for ∼16h prior to fixation with 2.5% glutaraldehyde in cacodylate buffer (100 mM sodium cacodylate with 2 mM CaCl_2_, pH 7.4) for 30 min. Fixed cells were incubated in DAB solution (1.3 mM DAB, 10mM H_2_O_2_ in PBS) for 10 min at room temperature and washed three times with PBS. Coordinates of DAB-stained cells were determined by brightfield microscopy. Then, cells were processed and imaged as described above except without post-staining.

### Analysis of mitochondrial morphology by fluorescence microscopy

To analyze mitochondrial morphology, untreated CRISPRi cells were grown directly to ∼60% confluency on glass-bottom dishes (Cellvis). For transient knockdowns, the indicated siRNAs were transfected at a final concentration of 20 nM with Lipofectamine RNAiMAX (ThermoFisher). The liposome/siRNA mixture was added directly to culture media for 24 h. Then, cells were passaged to glass bottom dishes for morphology analysis or culture dishes for whole cell lysate preparation to confirm knockdown efficiency by Western blotting. Cells were incubated for an additional 12-16h prior to analysis. Cells were treated 25nM Mitotracker Deep Red FM (ThermoFisher) for 30 minutes, washed once with growth media, and imaged with a Nikon Eclipse Ti inverted epifluorescence microscope equipped with a Hamamatsu Orca-Fusion sCMOS camera and a Nikon 100x 1.45-NA objective and acquired with Nikon Elements. All images were deconvolved using AutoQuant X3 (10 iterations, blind deconvolution, and low noise), and linear adjustments were made with Fiji. All data analysis/quantification was performed on nondeconvolved (raw) images using Fiji (see below). All z-series images were obtained using a 0.2-µm step size, and maximum projection images are shown. Samples were blinded prior to imaging and subsequent analysis and cells were manually categorized as fragmented, tubular, mildly enlarged, and severely enlarged or bulbous based on their mitochondrial morphology. Images were collected from three independent experiments, and approximately 100 cells were analyzed per experiment. Statistical comparison was performed between each sample by unpaired two-tailed t-test of the combination of mild and severe mitochondrial enlargement morphology categories.

### Mito-mKeima mitophagy assay

HeLa mito-mKeima cells were seeded at 200,000/well in a 6-well plate 12-16h prior to transient transfection with the indicated siRNAs (20 nM) with Lipofectamine RNAiMAX (ThermoFisher). The liposome/siRNA mixture was added to culture media for 24h, then 200,000 cells were passaged to 6-well plates and allowed to adhere 12-16h prior to a second round of transfection as above. After the second transfection, 75,000 cells were passaged into glass-bottom dishes and the remainder of cells were passaged to culture dishes for Western analysis to confirm knockdown at the end of the experiment. For untreated cells (Figure 6), cells were allowed to grow an additional 36h prior to imaging. To mimic hypoxia (Figure 7), cells were allowed to adhere for 12h and treated with CoCl_2_ (250 µM, Sigma) for 24 h prior to analysis. CoCl_2_-treated samples where BNIP3 and/or BNIP3L were depleted were simultaneously treated with 20 µM Q-VD-OPh (Apexbio) to prevent apoptotic cell death. Imaging was performed using a Zeiss Axio Observer microscope equipped with a Yokogawa CSU-W1 spinning disk module and a Photometrics Prime 95B sCMOS camera and a Zeiss 63x objective. Z-series images were acquired with a 0.2-µm step size. Detection of neutral mito-mKeima and acidified mito-mKeima were made using dual excitation with 473 nm (pH 7) and 561 nm (pH 4) lasers, respectively, and a 617/73 nm (73 nm bandpass filter centered at 617 nm) emission filter. Samples were blinded prior to analysis and the number of acidified mito-mKeima puncta per cell were manually counted in Fiji by examining single plane images throughout *z-*series of individual cells. Images were collected from three independent experiments, and 100 cells were analyzed per experiment. Data depicted graphically are the collective sum of data from all experiments.

## Acknowledgements

We thank Richard Youle (NIH) for generously providing HeLa mito-mKeima cells. We thank Mike Henne and Angelique Whitehurst (UTSW) for sharing plasmids and helpful discussions. The UTSW Flow Cytometry Core Facility provided experimental support for FACS sorting. The UTSW Proteomics Core Facility performed mass spectrometry analysis. The UTSW EM facility prepared samples for analysis and is supported by NIH 1S10OD021685-01A1. The UTSW Quantitative Light Microscopy Facility, which is supported in part by NIH P30CA142543, provided access to the Nikon SoRa microscope (purchased with NIH 1S10OD028630-01 to Kate Luby-Phelps) and deconvolution software. This work was supported by grants from the NIH to JF (R00HL133372 and R35GM137894) and the UTSW Endowed Scholars Program.

---

**Figure S1.**
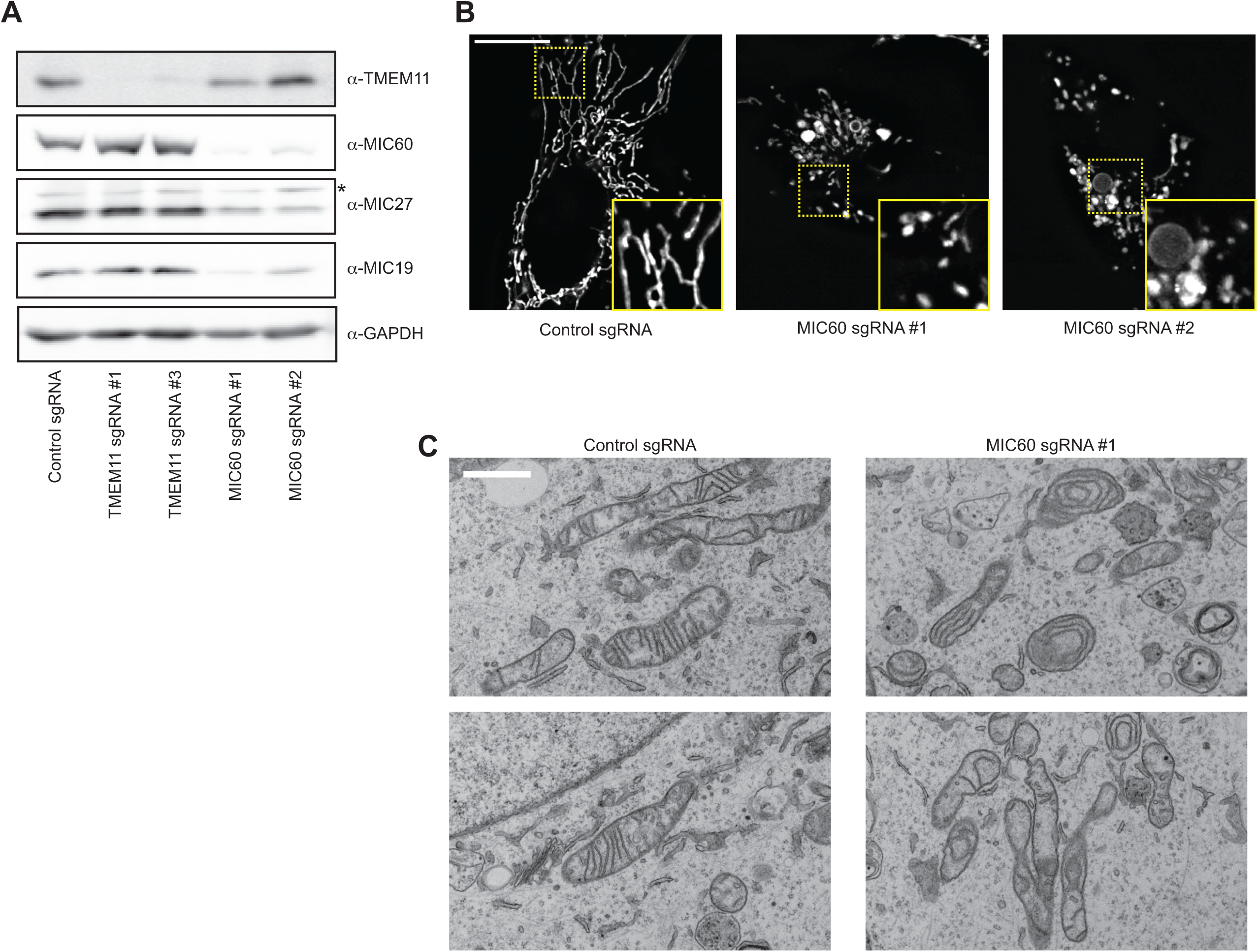
Depletion of the MICOS complex does not affect TMEM11 stability. **(A)** Western blot analysis of whole cell lysates from U2OS CRISPRi cells expressing scrambled control sgRNA or the indicated sgRNAs targeting TMEM11 or MIC60 and probed with the indicated antibodies. **(B)** Deconvolved maximum intensity projections of fluorescence microscopy images are shown of U2OS CRISPRi cells stably expressing the indicated sgRNAs and stained with Mitotracker Deep Red. Insets correspond to dotted boxes. Scale bar = 15 µm. **(D)** Representative electron micrographs of mitochondria from CRISPRi cells expressing control sgRNA (left) or sgRNA targeting MIC60. Scale bar = 1 µm.

**Figure S2.**
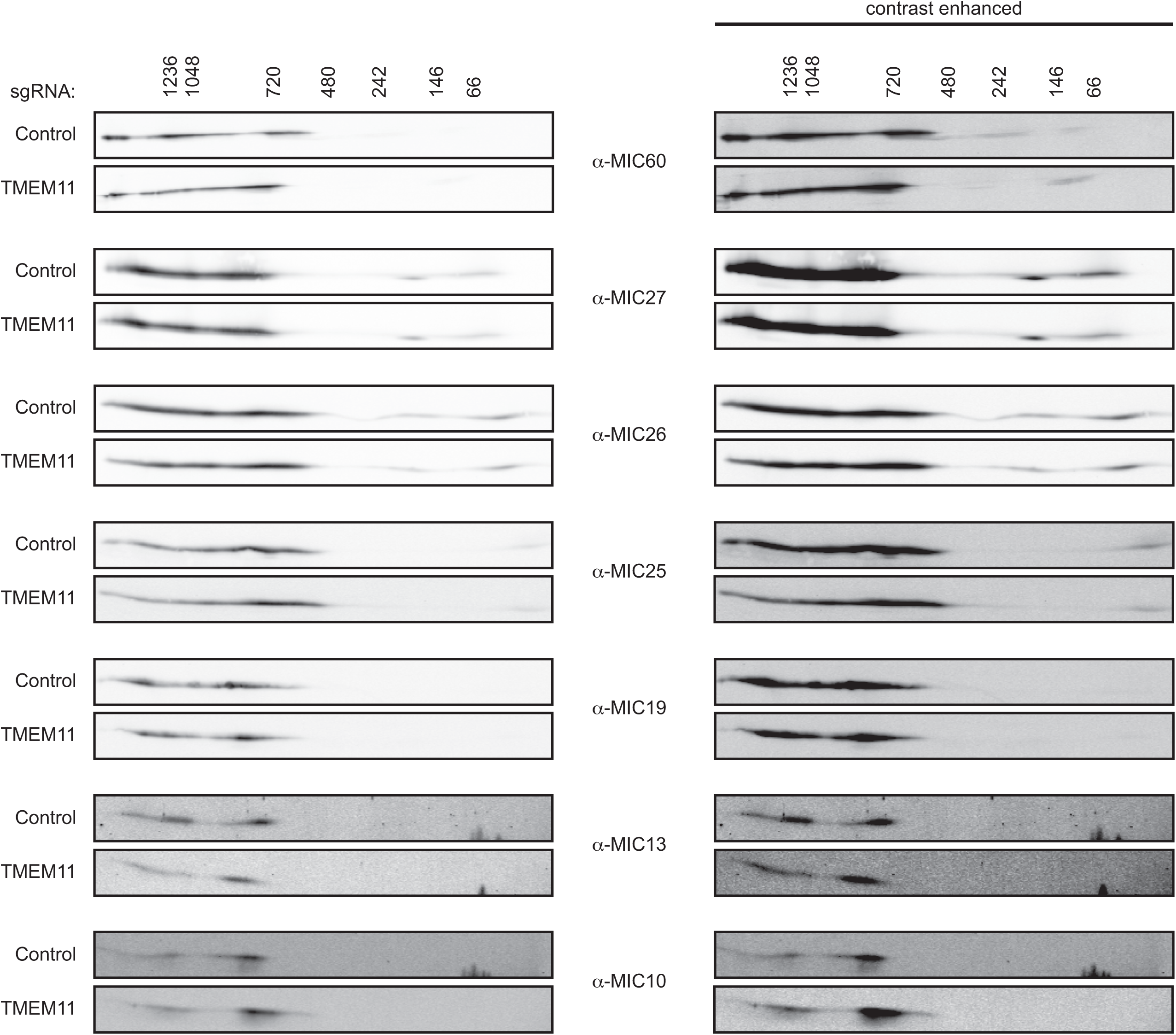
Lower molecular weight MICOS assemblies do not accumulate in the absence of TMEM11. 2D BN-PAGE and Western analysis of mitochondria isolated from U2OS CRISPRi cells expressing control or TMEM11-targeted sgRNAs and probed with the indicated MICOS antibodies. Images on left are linearly adjusted and redisplayed from Figure 2C. Images on right are contrast-enhanced to enable visualization of lower molecular weight MICOS assemblies.

**Figure S3.**
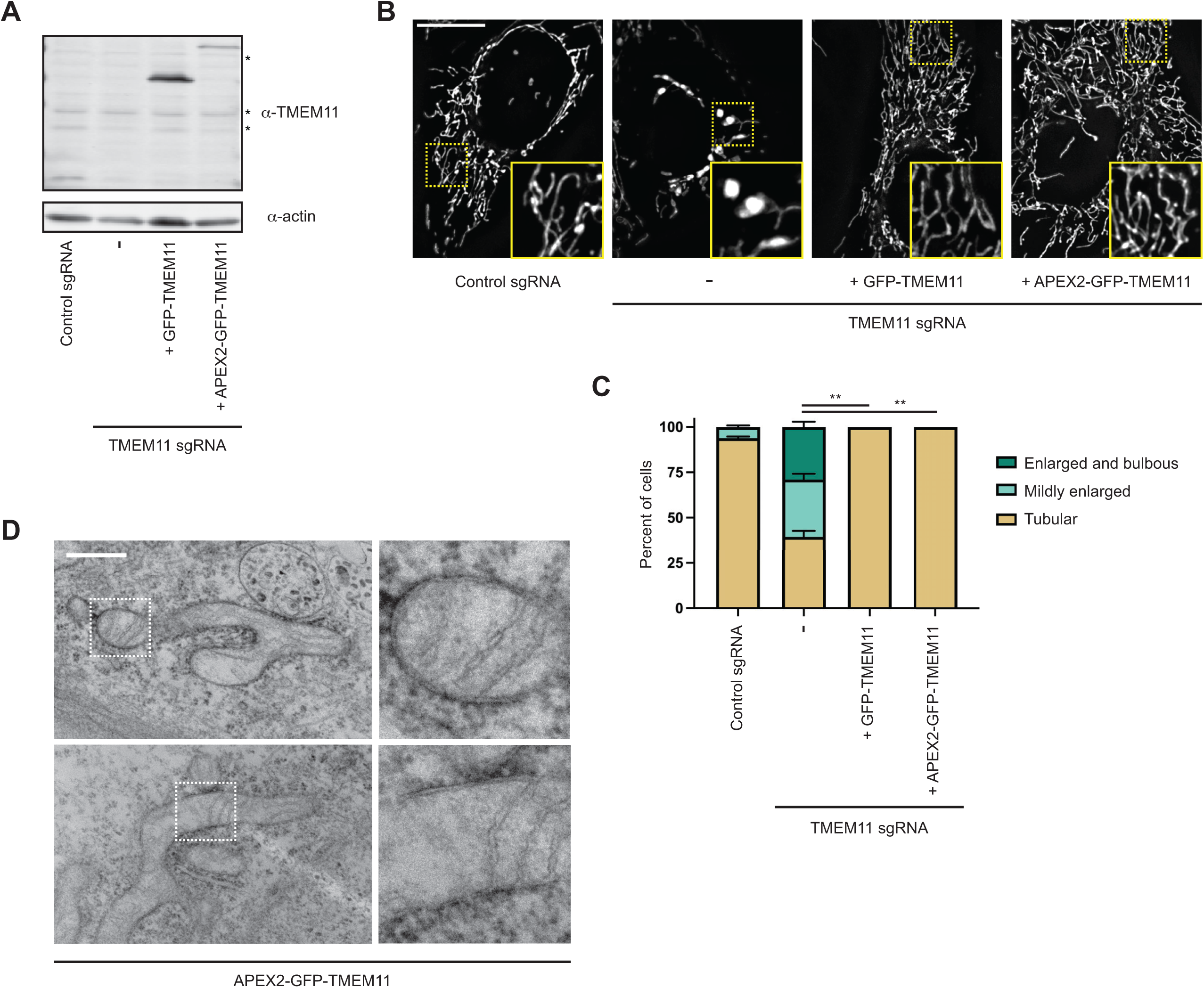
Mitochondrial morphology of TMEM11 CRISPRi cells can be rescued by reintroduction of N-terminally tagged TMEM11. **(A)** Western blot analysis with the indicated antibodies of whole cell lysates of CRISPRi cells expressing (left) control sgRNA or (right) TMEM11 sgRNA #3 cells that were lentivirally transduced with either GFP-TMEM11 or APEX2-GFP-TMEM11. Asterisks indicate cross-reacting bands. **(B)** Deconvolved fluorescence microscopy images are shown of cells as in (A) stained with Mitotracker Deep Red. Insets correspond to dotted boxes. Scale bar = 15 µm. **(C)** A graph of the categorization of mitochondrial morphology from cells as in (B). Data shown represent approximately 100 cells per condition in each of three independent experiments and bars indicate S.E.M. Asterisks (**p<0.01) represent unpaired two-tailed *t* test. **(D)** Additional examples of EM images from proximity labeling analysis of TMEM11 CRISPRi cells expressing APEX2-GFP-TMEM11. Enlargements (right) correspond to dotted boxes (left). Scale bar = 500 nm. See also Figure 3C.

## Notes

### Competing Interest Statement

The authors have declared no competing interest.

